# Crucial Roles of Carotenoids as Bacterial Endogenous Defense System for Bacterial Radioresistance of *Deinococcus radiodurans*

**DOI:** 10.1101/2021.05.26.445811

**Authors:** Qiao Yang

## Abstract

**Background:** The central dogma of radiation biology is that the cytotoxic and mutagenic effects of radiation are principally the result of biological macromolecules (DNA, RNA, polysaccharides, proteins and lipids) damage caused during the course of irradiation. Previous studies on the radioresistance of the extremely radiation-resistant red-carotenoid-pigmented bacterium *Deinococcus radiodurans* have been focused on DNA repair system. However, relative little is known about the biochemical basis of the extraordinary recovery capacity of *D. radiodurans* for ionizing radiation and can include hundreds of DNA double-strand breaks.

**Methodology/Principal Findings:** Here we identified a novel carotenoid in *D*. *radiodurans* as ((all-E)-1’-hydroxy-3’,4’-didehydro-1’,2’,2,3-quahydro-β,ψ-carotene-4-one-1) and characterized the intracellular distribution of carotenoids. Following exposure to ionizing radiation, the wild-type parent of *D. radiodurans* survived much higher compared with radiation-sensitive colorless mutants screened by ^60^Co irradiation and by knockout of *crtB* or *crtI* genes for the carotenoid biosynthesis. Electron paramagnetic resonance spectroscopy analysis showed that two carotenoid can effectively scavenge radicals including superoxide anion (O_2_^**.−**^) and hydroxyl radical (^.^OH). Consistent results showing the substantial antioxidant abilities of the two carotenoids were obtained by UV-induction of bacteriophage λ method in vivo, and the detectable reaction potentials of two carotenoids with nitric oxide (NO) evaluated by ultraviolet-visible near-infrared ray (UV/vis/NIR) spectra assay.

**Conclusions/Significance:** Our findings support the idea that the degree of bacterial radioresistance of *D. radiodurans* is partly related to the levels of carotenoids in the cells. Carotenoids, as an important member of the structural and non-structural components of the cells, are endogenous scavengers of reactive oxygen species and can effectively quench radicals generated during irradiation. The current work suggests that carotenoids play a previously unrecognized protection role in the endogenous defense System (EDS) that facilitates the recovery from radiation injury. And we propose a complete model to elucidate the mechanisms of extreme radioresistance in this remarkable bacterium.

## Introduction

*Deinococcus radiodurans* is a red-pigmented, non-spore-forming, nonmotile spherical bacterium that was originally identified as a contaminant of canned meat irradiated at 4,000 grays (Gy) of γ radiation, a dose that 250-times higher than that used to kill *Escherichia coli* [1]. *D*. *radiodurans* is the type species in the genus *Deinococcus* that exhibit remarkable resistance to the lethal and mutagenic effects of a variety of genotoxic damaging agents and conditions, particularly the effects of ionizing radiation (IR). Depending upon conditions for its growth, this bacterium is capable of surviving a startling 15,000 Gy (1.5 million rads) of IR, a thousand times more than any other known organism including humans, and it shatters the bacterium’s chromosomes into hundreds of fragments. Cool or freeze the microbe, and it may survive 30,000 Gy of IR [2–4].

Relatively little is known about the biochemical basis of the capacity of *D. radiodurans* to endure the genetic insult that results from exposure to IR. IR can include hundreds of DNA double-strand breaks (DSBs) – the most difficult to repair and therefore the most lethal form of DNA damage. Studies to uncover the extraordinary radioresistance are mostly focused on the highly efficient DNA repair system [3–16]. Evidence is accumulating for both passive and enzymatic contributions to bacterial radioresistance of *D. radiodurans*. However, the molecular mechanisms responsible for this specie’s extraordinary resilience remain poorly understood. Further study based on alternative genetic and biochemical approaches should help to gain a better understanding of the radioresistance mechanisms of *D. radiodurans* [17–18].

Carotenoids represent one of the most widely distributed and structurally diverse classes of natural pigments in plants, microorganisms, animals and humans, producing colors ranging from light yellow through orange to deep red [19,20]. Presently, at least 600 different carotenoids are naturally synthesized, performing important functions in photosynthesis, nutrition and protection from oxidative damage. Some carotenoids exhibit provitamin A activity, some might function as significant dietary antioxidants. Other beneficial effects of carotenoids are currently under investigation including inhibition of carcinogenesis, enhancement of the immune response and prevention of cardiovascular disease [20,21]. *D*. *radiodurans* R1 is red-carotene-pigmented and the main carotenoid has been identified as deinoxanthin [(all-E)-2,1’-dihydroxy-3’,4’-didehydro-1’,2’-dihydro-β,ψ-carotene-4-one-1] by Lemee et al [22] and Saito et al [23].

Currently, the biological roles of carotenoids for the bacterial radioresistance of *D. radiodurans* remain unclear. In this report, we demonstrate direct evidence of the chemical structure of a novel carotenoid in *D. radiodurans* R1, and investigations concerning the intracellular distribution, antioxidant and radical scavenging functions of the two major carotenoids. This study reveals that carotenoids, as an important member of the structural and non-structural components of the cells, play a crucial role in the systematically described endogenous defense System (EDS) for radioresistance of *D. radiodurans*. Based on accumulative reports, we proposed a complete model for the mechanisms of extreme radioresistance of the bacterium.

## RESULTS

### Constitutions of *D. radiodurans* carotenoids

The carotenoids extracted from *D. radiodurans* cultures were separated and analyzed by high-pressure liquid chromatography (HPLC) monitored by a diode-array detector set at 470 nm. In comparison to the results reported previously [22,23], the other main peak (peak 3) except for the previously identified deinoxanthin (DE, peak 1) was found in HPLC chromatogram (Figure 1), and the quantitative analysis demonstrated that the relative amount of the two peaks accounted for 57% and 28% of the total carotenoids in *D. radiodurans* R1, respectively. The fine structures of UV-Vis spectra of two main peaks have almost the same maxima at 451, 480 and 505 nm indicating that the conjugated polyene system contained twelve double bonds [22,24,25]. Reduction of this compound by NaBH_4_ gave a carotenoid with a UV/Vis spectrum of a pronounced fine structure and maxima at 457, 475 and 506 nm in ethanol. This confirmed the presence of a carbonyl group conjugated to the polyene chain [22,24,25]. APCI-MS/MS analysis showed the molecular ion [M+H]^+^ at *m/z* 567.30, corresponding to a molecular formula C_40_H_56_O_2_. Combined with the data from UV-Vis, MS/MS spectra as well as chemical properties tests, this novel carotenoid was identified as (all-E)-1’-hydroxy-3’,4’-didehydro-1’,2’,2,3-quahydro-β,ψ-carotene-4-one-1. This carotenoid has not yet been reported in nature, so we proposed the name dehydroxydihydro-deinoxanthin (DD-DE). Peak 2 in HPLC chromatogram was the trans-isomer of deinoxanthin based on the mass spectra and UV-vis spectra data (Figure 1). The characterization of this novel carotenoid confirmed and extended the previous evidence [22,23] that this bacterium contains a variety of carotenoids of unconventional structures.

**Figure 1.**
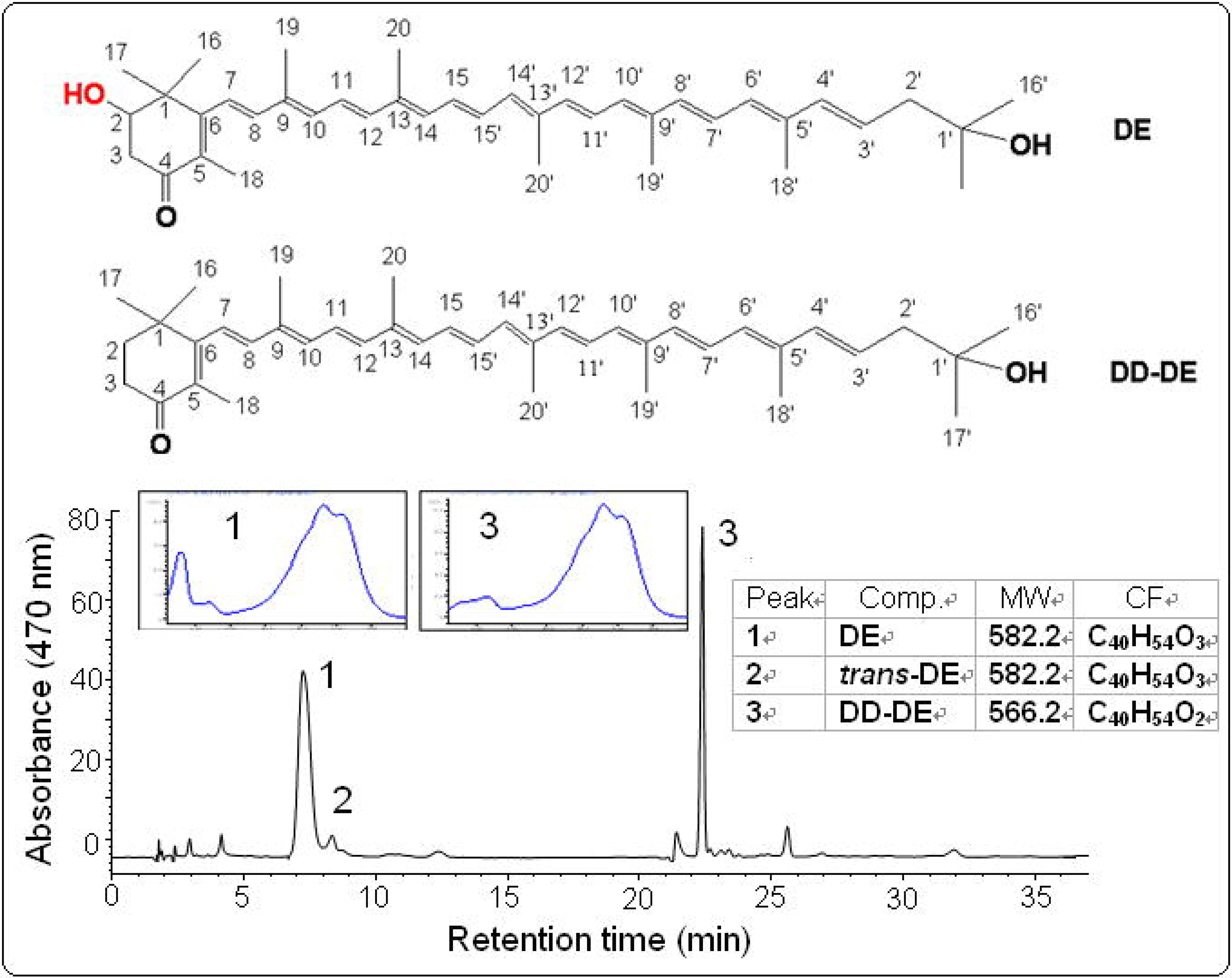
HPLC profile and constitutions of *D. radiodurans* carotenoids. A total of 250 mg of wet cultures at exponential phase were extracted and then analyzed by reversed phase HPLC recorded at 470 nm with a diode-array detector. Top, chemical structure of deinoxanthin. Low, chemical structure of dehydroxydihydro-deinoxanthin, Comp, Compound; MW, molecular weight; CF, chemical formula.

### Intracellular distribution of *D. radiodurans* carotenoids

Owing to the indirect effect by the induced reactive oxygen species (ROS) and direct interaction between γ-photons and macromolecules during irradiation exhibit spatial distribution effect, the intracellular protection of carotenoids may not highly dependent on their absolute concentrations in the cell. In this context, we examined the dynamic accumulation of *D. radiodurans* carotenoids using HPLC analysis [22,23] (Figure 2), and the intracellular distribution of carotenoids using pretreated the cells with lytic enzymes and followed by density gradient centrifugation and HPLC analysis (Figure S1) [44]. The inner layer of the cellular envelope is the plasma membrane common to all cells. The lipids of the plasma membrane are atypical, containing unique fatty acids. Unlike phosphoglycolipids in other organisms, these lipids contain alkylamines. Cell walls of *D. radiodurans* are consisted of at least four unusual layers that are separated from each other, including an outermost network structure which can be removed by trypsin, a fragile soft layer containing hexagonal packed intermediate (HPI) layer consisting of tightly-packed hexagonal proteins, and a rigid layer riddled with perforations, and so dubbed “the holey layer” with complex outer membrane lipids, a compartmentalized layer whose structure is not known yet, and a thick typical peptidoglycan layer atop the plasma membrane containing the amino acid omithine [26]. The HPI layer was less dense, pink in color, and it contained carotenoids [26–29]. HPLC analysis of the sediment of cell wall of *D. radiodurans* culture demonstrated that the majority of carotenoids appeared in membrane. Carotenoids in cell walls may be associated with lipids or proteins in the hexagonally packed subunits [27,29]. While, a little carotenoids was also detected in the cytoplasm (Figure S1).

**Figure 2.**
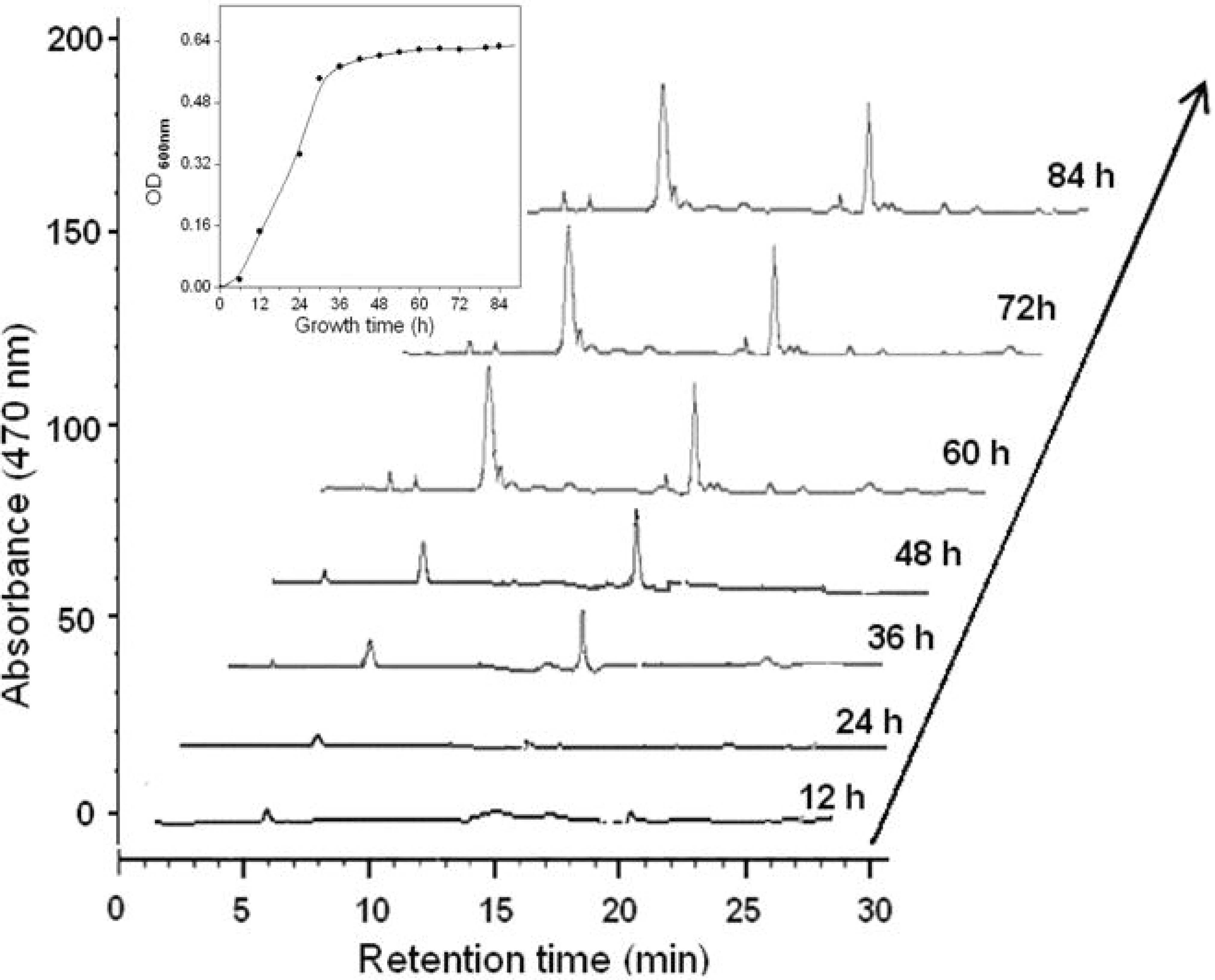
HPLC profile for the dynamic accumulation of *D. radiodurans* carotenoids during different growth phases.

### Colorless mutants of *D. radiodurans* were highly sensitive to IR

To elucidate the biological functions of carotenoids for radioresistance of *D. radiodurans*, we screened colorless mutants of *D. radiodurans* by ^60^Co irradiation with the total dose of 8 kGy. Approximately 2,000 colonies were screened; 15% appeared white duo to inactivation of the desaturase in carotenoids biosynthetic pathway. HPLC analysis showed that the two major carotenoids in wild-type strain of *D. radiodurans* R1 did not appear in the cultures of the colorless mutants, indicating disruption of the key synthase genes for carotenoids biosynthesis occurred in these mutants (Figure S2). To determine if loss of carotenoids affected the radioresistant capacities of these mutants, the recovery survival of the cultures for the wild and colorless mutants subjected to different dose of γ-irradiation (maximum to 12 kGy) were followed. Aliquots of exponential phase cultures were exposed to γ-irradiation. The wild-type strain exhibited the greatest resistance to irradiation with a D_10_ dose of 9,123 Gy (Pane A in Figure 3), whereas the colorless mutants showed a significant decrease of radioresistance at doses higher than 2,000 Gy (*P* < 0.02) and its D_10_ was 5,065 Gy. This assay provided direct evidence that the pigmented bacteria of *D. radiodurans* R1 were more resistant to the elevated dose of γ-irradiation than the colorless ones, while the colorless bacteria were as much as 100-fold more sensitive to the lethal effects of IR and their sensitivity was dose-dependent. Moreover, it should be noted that the colorless mutants cultures supplemented with the carotenoid of deinoxanthin (DE) from *D. radioduran* is as much as 8-fold more resistant to the lethal effects of γ-irradiation than the colorless without adding any carotenoid. So, our observation supports the conclusion that carotenoids play direct protective role for the cell survival subjected to γ-irradiation. These results are consistent with the ones for evaluation of the decolorization and decrease in light absorbance (470 nm) of *D. radiodurans* carotenoids subjected to different dose of γ-irradiation (Pane B in Figure 3). Furthermore, we previously constructed colorless mutants of *D. radiodurans* by knockout of two key genes (*crtB* and *crtI*) in the carotenoid biosynthetic pathway, and compared the cell survivals of two resulting colorless mutants with the parental strain [30]. We found similar evidence that colorless mutants were more sensitive to IR, UV, and hydrogen peroxide, but not to mitomycin-C compared to their wild-type parent [30]. Thus, these results support our model that carotenoids, largely present in parental strain, could act as endogenous radical scavengers and effectively quench ROS generated within the cells during irradiation, and therefore they are crucial part of EDS of *D. radiodurans*.

**Figure 3.**
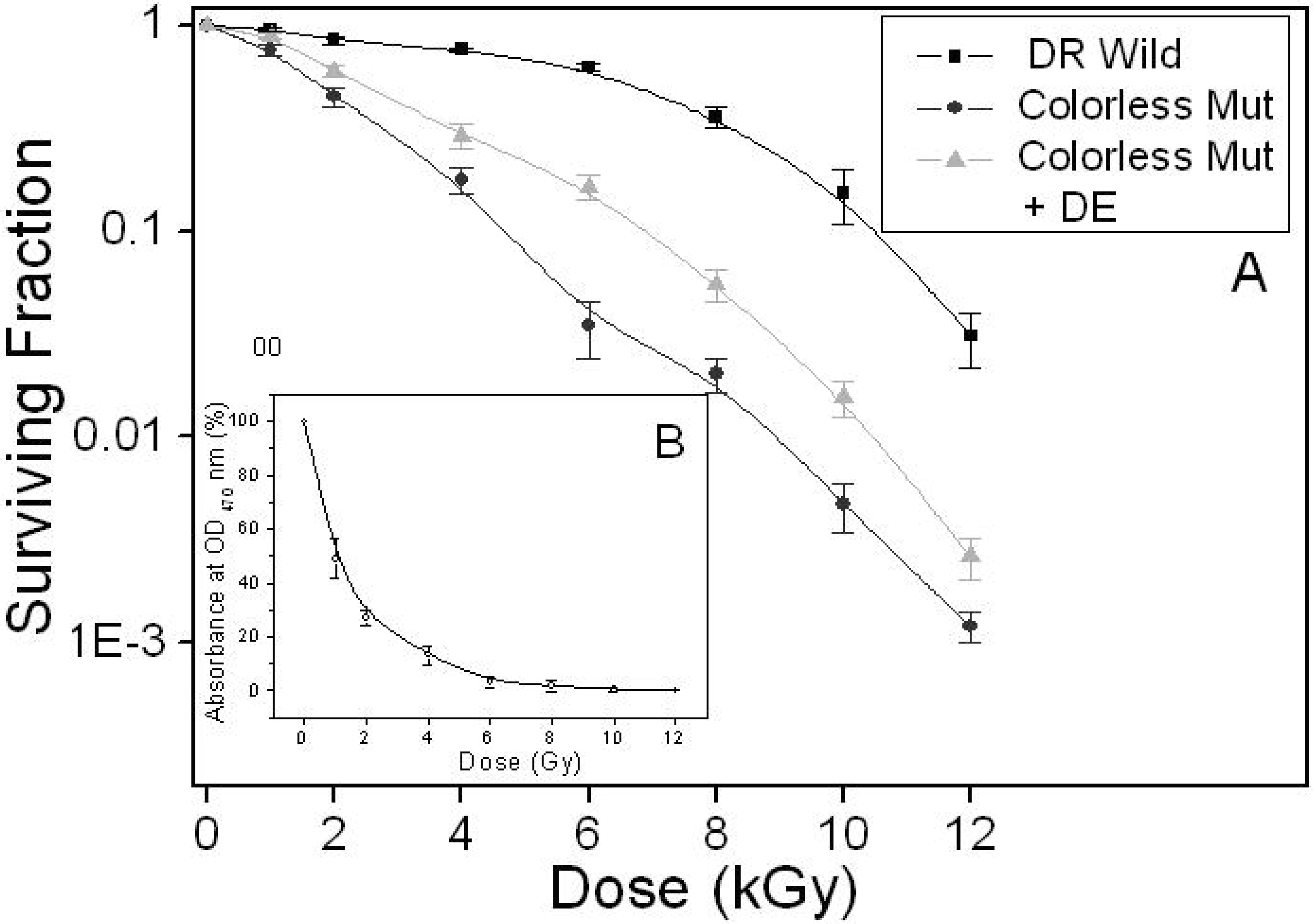
IR resistance of parental and mutants of *D. radiodurans.* Pane A presents the survival curves of parental strain of *D. radiodurans* R1 (■), colorless mutants (●) and colorless mutant cultures added with deinoxanthin (DE, 0.1 mM) (▲) subjected to γ-irradiation. Pane B presents the decolorization and decrease in light absorbance (470 nm) of the methanol extract from *D. radiodurans* R1cultures subjected to γ-irradiation. Values are the mean ± standard deviation of three independence experiments; *n* = 6.

### Two *D. radiodurans* carotenoids demonstrated substantial antioxidant and free radical scavenging abilities

To demonstrate a mechanistic link between antioxidant and free radical scavenging abilities of carotenoid and the capacity of *D. radiodurans* to endure the insults that results from exposure to IR. We examined antioxidant and free radical scavenging abilities of two *D. radiodurans* carotenoids both in vivo and in vitro. Using assays based on EPR which is the most direct and effective technique for the detection of free radical, and spin-trapping technique using 5,5-dimethyl-1-pyrroline-N-oxide (DMPO) [31,32], we tested whether or not the two *D. radiodurans* carotenoids can react with the main free radicals, O_2_^**.−**^ and ^.^OH, generated during irradiation. Both carotenoids were found to react strongly with O_2_^**.−**^ and ^.^OH and the effects were both in dose-dependent manners (Figure 4). By using a modified UV/vis/NIR spectra assay [33] (Figure 5), we also tested the reaction potential of the two carotenoids with nitric oxide (NO) which is one of ROS and a ubiquitous gaseous signal involved in the regulation of so diverse physiological/pathophysiological mechanisms in both mammalian and nonmammalian species [34]. Some of the cytotoxic effects of NO have been proposed to be mediated by it reaction with O_2_^**.−**^ to produce peroxynitite (ONOO^−^) - a very powerful oxidant and nitrating agent.

**Figure 4.**
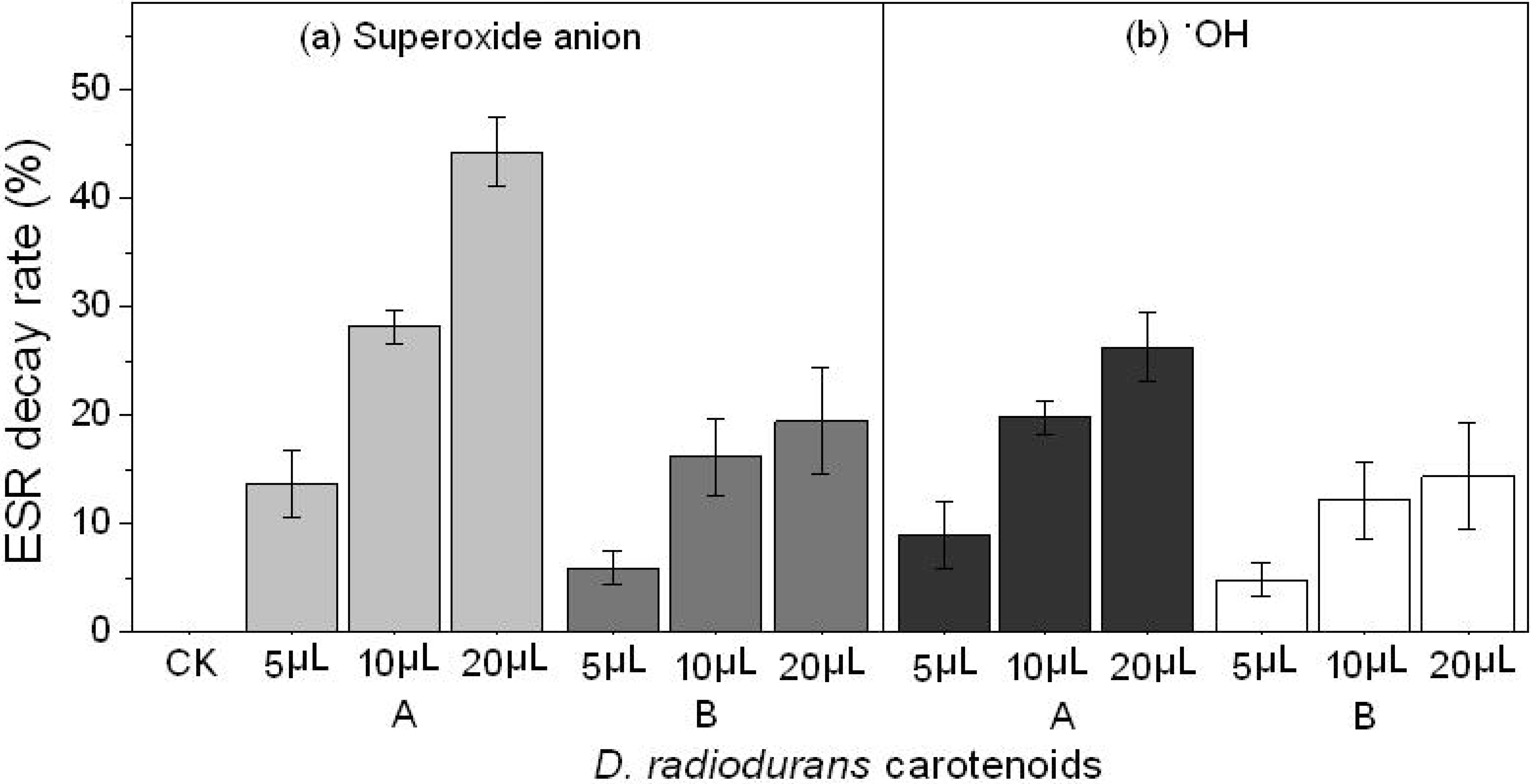
EPR and spin-trapping technique assay showing the scavenging effects of *D. radiodurans* carotenoids on O_2_^**.−**^ and ^.^OH. (A)DE; (B) DD-DE. Values are the mean ± standard deviation of three independence experiments; *n* = 6. 5,5-dimethyl-1-pyrroline-N-oxide (DMPO, 10 mM) was used as the spin-trapper.

**Figure 5.**
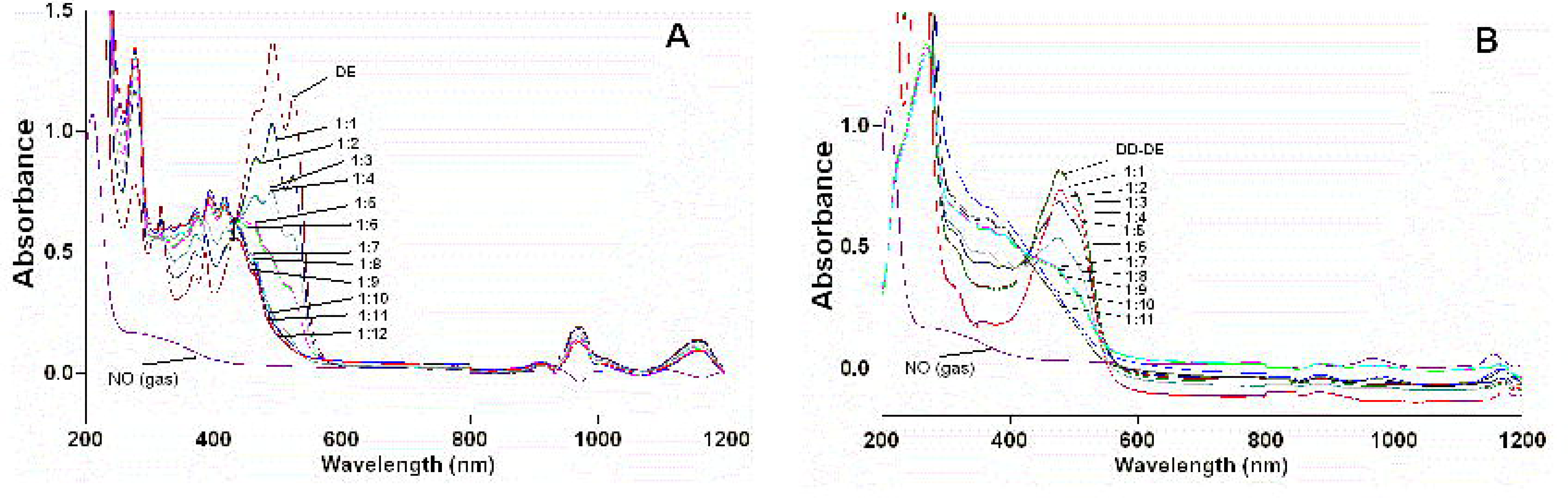
UV/vis/NIR spectra assay showing the bleaching of two *D. radiodurans* carotenoids by increasing the concentration of nitric oxide (NO). (A) DE; (B) DD-DE. Two carotenoids solution in methanol was rendered anaeroically, and small volume additions of a saturated NO solution in PBS buffer were added to give the indicated (carotenoid:NO). The spectra were recorded with a double beam Virian UV-5000 UV-VIS-NIR spectrophotometer (190-1200 nm). For the final spectrum NO gas was bubbled through the solution.

Carotenoids are widespread in the cells, particularly under conditions of oxidative stress initiated by some factors such as IR. Under such conditions, membrane and LDL (low density lipoprotein) peroxidation is accompanied by conjugated diene formation in the fatty acyl chains [35]. Therefore, besides its well known metabolic activities and as a significant widespread structural constituent, their effective antioxidant and free radical quenching capacity of the carotenoids in *D. radiodurans* assign them with an important protection role in EDS against oxidative stress initiated by IR. By using UIB method [36], we also established in vivo antioxidant and free radical scavenging ability of two *D. radiodurans* carotenoids. the results showed that two carotenoids also exhibited substantial inhibitory effects on UIB system through fighting free radicals generated during the induction process (Figure 6a), consisting with our studies that the two carotenoids showed demonstrable protective effects on *E. coli* subjected to UV irradiation (total dose 150 J/m^2^) (Figure 6b) and on the colorless *D. radiodurans* mutants (total dose 700 J/m^2^) (Figure 6c). The maximum inhibitory rates on UIB system of two carotenoids (DE and DD-DE) were 74% and 28%, respectively. The differences in the free radical scavenging abilities of carotenoids were directly related with their length of the conjugated double bonds and the functional group substitutions of the β-ionone ring [37,38].

**Figure 6.**
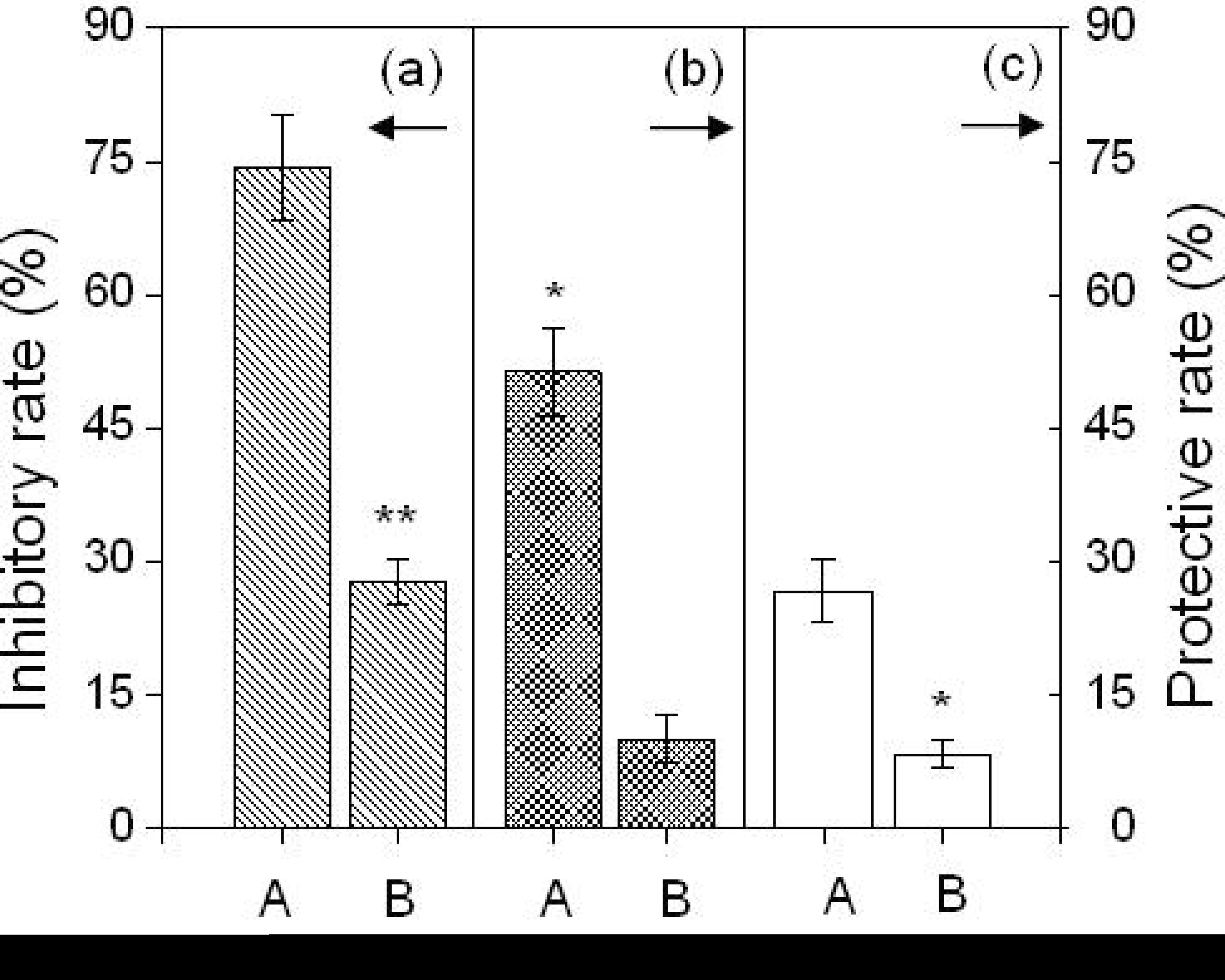
UIB method in vivo evaluating the antioxidant effects of *D. radiodurans* carotenoids (Pane a). The cell density of *E. coli* K12 (λ^+^) and irradiation time were fixed as 10^8^ cells/ml and 30s (60.1 J/m^2^), respectively. The concentrations of all carotenoids were kept as the same at optical density at 470 nm (OD_470nm_)_._ Pane b represents the protective effects of *D. radiodurans* carotenoids on *E. coli* subjected to UV radiation (150 J/m^2^). Pane c represents the protective effects of the carotenoids on the colorless mutant strains of *D. radiodurans* subject to UV irradiation (700 J/m^2^). (A) DE; (B) DD-DE. Values are the mean ± standard deviation of three independence experiments; *n* = 6. *P*<0.05; ** *P*<0.02.

## DISCUSSION

Ionizing radiation can induce free radicals in chemical and biological systems. Ions and free radicals produced pass through matter, rapidly react and modify molecules on a nanosecond timescale, hence cause a series of considerable biological damage, including chemically altering the major classes of biological macromolecules: polysaccharides, proteins and lipids and nucleic acids (Figure 7) [6]. Typically during irradiation, ~80% of DNA damage is caused indirectly by ROS induced during irradiation, and the remaining ~20% by direct interaction between γ-photons and DNA [7,39]. Highly reactive hydroxyl radical (^.^OH), a primary product of the radiolysis of water, is extremely toxic. In the presence of O_2_ it can generate other ROS, including superoxide ions (O_2_^.–^) which results from the rapid reaction between solvated electrons (e^−^_aq_) and O_2_ and hydrogen peroxide (H_2_O_2_) [39,40]. H_2_O_2_ is relatively stable and diffusible, but it is decomposed to ^.^OH through the Fenton and Haber-Weiss reactions in the presence of free Fe(II) in the cell (Figure 7). Among all the effects of radiation, genome damage probably has the greatest impact on cell viability [6]. IR generates multiple types of DNA damage and as much of the damage results from the oxidative action of hydroxyl radical. In DNA, DSB is regarded as the most difficult to repair and therefore the most lethal form of DNA damage [7,15]. DSB can result in significant loss of genetic information and, if not repaired, will prevent replication of the prokaryotic genome. Therefore, how an organism deals with DSB lies at the heart of that organism’s capacity to survive radiation exposure [6]. However, this emphasis has been challenged by recent report indicating that protein, rather than DNA, is the principal target of the biological action during irradiation in sensitive bacteria [10]. The consequences of a series of considerable oxidative damage to all major classes of biological macromolecules including RNA, polysaccharides, lipids and proteins, not the only of DNA, caused by IR is a continual problem that cells must guard against to survive. Although there is a relatively conventional set of DNA repair systems but with far greater efficiency existing in this remarkable bacterium, it alone can not support the extraordinary resistance to the lethal and mutagenic effects induced by lots of bacterial damaging agents and conditions including IR, desiccation, UV radiation, or electrophilic mutagens [5–16]. On the other hand, it’s reasonable to speculate that the repair systems accounting for radioresistance of *D. radiodurans* may not limited to DNA repair, proteins and lipid repair systems and even cell ultrastructural reconstruction may also be involved (Figure 7).

**Figure 7.**
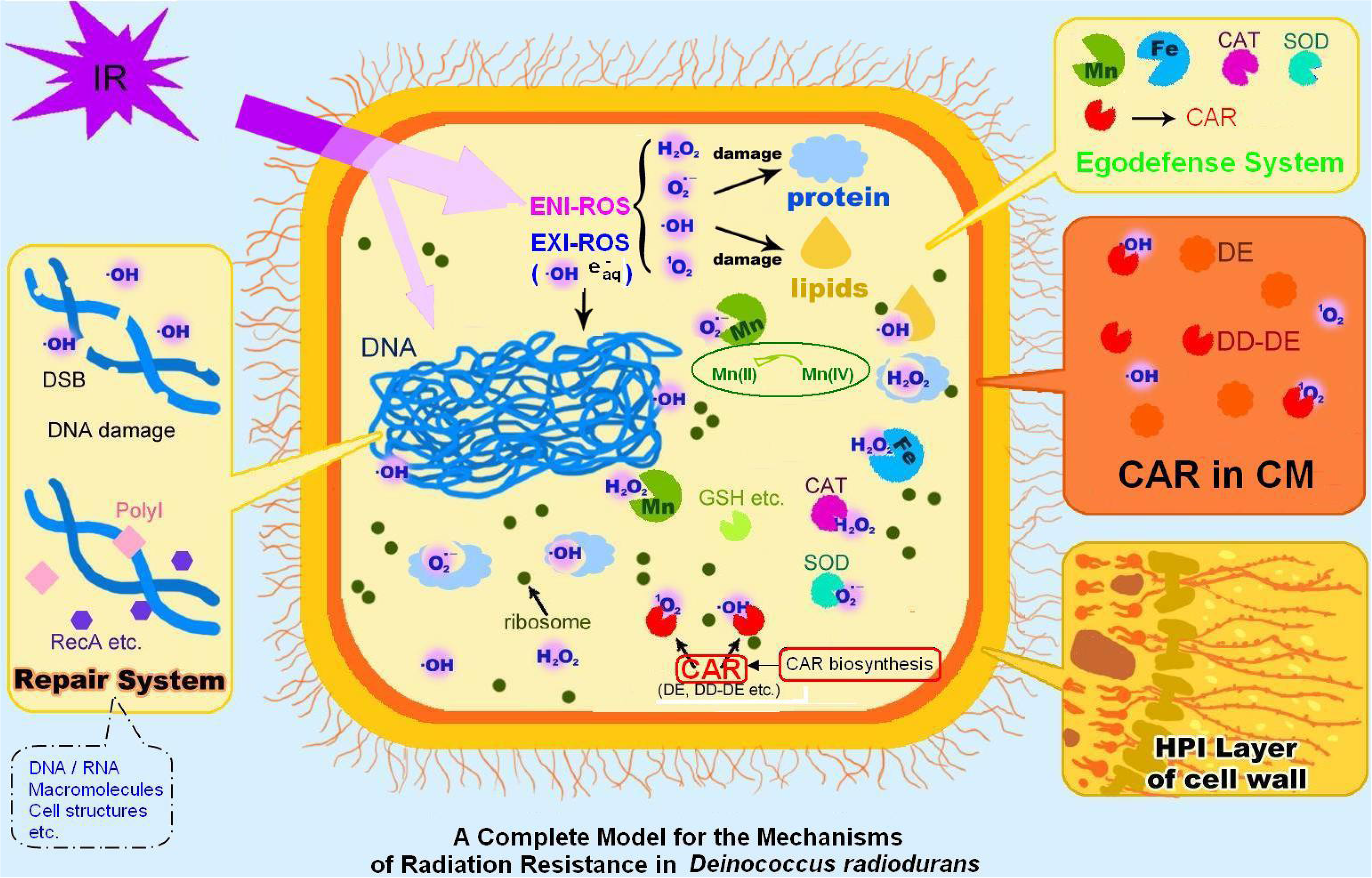
Proposed complete model for radioresistance mechanism of *Deinococcus radiodurans.* The locations of a cell are color-coded as follows: khaki, cell wall; orange, membrane; ivory-white, cytoplasm. Abbreviations: CAR, carotenoids; CAT, catalase; CM, cell membrane; DE, deinoxanthin; DD-DE, dehydroxydihydro-deinoxanthin; DSB, double-strand break; e^−^_aq_, hydrated electrons; ENI, endogenously induced; EXI, exogenously induced; H_2_O_2_, hydrogen peroxide; HPI, hexagonally packed intermediate; IR, ionizing radiation; NO, nitric oxide; O_2_^**.-**^, superoxide anino/radical; ^.^OH, hydroxyl radical; PQQ, pyrroloquinoline quinine; ROS, reactive oxygen species; SOD, superoxide dismutase.

Cellular response to γ-irradiation-induced oxidative stress entails complex mRNA and protein abundance changes, which translate into physiological adjustments to maintain homeostasis as well as to repair and minimize damage to cellular components [41]. In radioresistance research, the most interest has been on the offensive side, for example, the roles of ionization effects on biological macromolecules, and oxygen radical intermediates. Little has been focused on the defensive side, which is responsible for the attenuation and resolution of oxidative stress. As key features of DNA-centric hypotheses of extreme resistance have grown weaker, the study of endogenous defense mechanisms/systems in the cells should be given our exclusive attention, mostly because of their relative biological complexity. Microbes have evolved strategies to avoid foreign attacking that interrupt the homeostasis system at many levels, and one of the most sophisticated is mounted by some cellular structures which have multiple mechanisms for self-protection.

The intriguing array of captivating features that *D. radiodurans* offers have been studied with great detail over decades. At present, two systems, endogenous defense and repair systems, are believed to be responsible for its remarkable radioresistant ability. The repair system in *D. radiodurans* has been extensively studied [5–9,15,16]. However, the endogenous defense gained little attention. EDS represents the active and inherent strategy by which bacteria employ to protect themselves against environmental stress such as IR [42,43]. It could be generally classified into two types: structural and non-structural components. The structural components includes the complex and unusual HPI layer in the periplasm of *D*. *radiodurans* [26,27], ringlike nucleoids organization and the high degree of genome condensation etc. The mechanism of radioresistance of *D. radiodurans* is believed largely ascribed to highly efficient systems of DNA repair [3,5–7]. However, cells of *D. radiodurans* appear to resist radiation-induced membrane damage to a greater degree than do cells of radiation-sensitive bacteria [44,45], suggesting that structural membrane components may be involved in one aspect of EDS in radioprotection. The contribution of the cell structures to resistance to ionizing and UV radiation is not clear, but it has been suggested that the bacterial cell wall and pigmentation may contribute to the overall DNA damage (14). Moreover, Daly et al. [4] has demonstrated that *D. radiodurans* is still comparatively resistant to IR when they inhibit the Mn redox (D_10_ = 6,000 Gy) or even completely deplete Mn (D_10_ = 2500 Gy), these data strongly indicate that other factors (like carotenoids in EDS) must exist and contribute to the residual radioresistant ability. Like all gram-positive bacteria, *D. radiodurans* has evolved to equip or carry usual cell structures such as HPI layer which resists penetration into the cell membrane by the membrane-attack complex, that means this bacterium cannot be easily damaged by oxidative stress. This forms a first level of bacterial endogenous defense (Figure 7). Currently, little is still known about the protection functions of the structural components of either the membranes or of the relatively complex cell envelope layers of *D. radiodurans* for its radiation resistance [46–48]. Chemical characterization of these constituents will, therefore, shed light on understanding the roles of such boundary structural layers in radioresistance. Carotenoids are constituents of the protein-lipid complex, low density lipoprotein (LDL) and the in vivo cholesterol transport particle. Moreover, conjugated system of double bonds, of which carotenoids (DE and DD-DE) are only one, are widespread particularly under conditions of oxidative stress. Under such conditions membrane and LDL peroxidation is accompanied by conjugated diene formation in the fatty acyl chains. Moreover, nitric oxide (NO) is more stable in the hydrophobic phase presented by lipids and thus conjugated diene-NO reactions would be favoured. Non-structural components refer to antioxidant enzymes and antioxidants/free radial scavengers. During radiation, most cellular damage are caused indirectly by ROS, which are highly reactive and can attack major classes of biological macromolecules, resulting in various damages on the cell membrane structure, enzymic system or genetic material. The growth and duplication of bacteria will be greatly affected as the IR doses increased. The non-structural components of EDS are dedicated to protect those macromolecules and cell structures from IR-induced detrimental effects by directly scavenging or quenching ROS. The non-structural components also include Cu/Zn superoxide dismutase (Cu/Zn SOD) to scavenge O_2_^**.-**^, catalase (CAT) to scavenge H_2_O_2_, peroxidase (Px) and glutathione peroxidase (GPx) to scavenge hydroperoxides, peroxiredoxin (Prxs) [49,50] to carry out the cellular detoxification of H_2_O_2_ and organic hydroperoxides (OHPs) via a thioredoxin-recycling process, and coenzyme pyrroloquinoline-quinone (PQQ) protein [51] etc. The triggered PQQ enhanced expression/activity of antioxidant enzymes has been proposed as one of the mechanisms underlying the impressive oxidative stress tolerance of this superbug [51]. While the unusual layering of *D. radiodurans* may well constitute a lead vest, the endurance of the organism may be credited to a combination of special unidentified antioxidants like PQQ existing in EDS for its egregious resistance. The antioxidants/free radial scavengers of non-structural components in *D. radiodurans* include phospholipids [44], natural lipid (like vitamin MK8) [52], and Mn(II) [4] etc. Daly et al [4] has recently reported that intracellular Mn(II) can be protective for proteins, but not DNA by quenching ROS and related free radicals produced during irradiation. In face, different free radical scavengers are responsible for quenching specific species of radicals or ROS. Although their structural requirements are well established, the molecular functions of the structural components are ill-defined. Despite all the information currently known, EDS is rather complex and many factors may not be appreciated yet and await further investigations.

It should be noted that membrane phospholipids of *D. radiodurans* have been implicated as functioning in membrane transport of ions and solutes, which would be especially important in regulating and reducing ion leakage during radiation [51]. It is interesting to speculate that either the complex wall structure of *D. radiodurans* [44] or its unconventional membrane phospholipids components may be involved in protecting *D. radiodurans* from the lethal and mutagenic effects of radiation on membrane structure and biological function. In other systems, radiation sensitivity has been shown to vary with fatty acid composition [53]. Moreover, specific classes of lipids with antioxidant properties [54,55] may be important in moderating radiation-induced membrane damage. In the present study, intracellular distribution of *D. radiodurans* carotenoids in cells demonstrated that majority of the carotenoids appeared in cell membrane (Figure 7 and S1), and the length of the carotenoid molecule corresponds approximately to the width of the membrane lipid bilayer [38]. With an orientation perpendicular to the membrane plane, the conjugated system of double bonds with functional carbonyl and hydroxyl groups could trap free radicals at any depth in the membrane, and provide immediate protection from ROS, and facilitate removal of fatal damages from cells exposed to IR [56]. On the other hand, the cell walls also contained two major carotenoids. As the outmost and first line of defense against irradiation in the cells, structural carotenoids in cell walls may perform the similar active and protective functions as they do in membrane and cell cytoplasm (Figure 7), and making it differenced with the predominant protection of Mn(II) to the cytosol previously reported by Daly et al. [10].

The importance of repair system should also be noted especially considering about 20% of chromosomal damages are caused by direct interaction between γ-photos and DNA. And the system is also crucial for the repair of macromolecular damages of which this generation exceeded the protection capacity of the EDS. We believe the repair system in *D. radiodurans* supervises not only DNA, but also proteins and lipids, and maybe the damaged cell structure. endogenous defense is an active and effective strategy for bacterial adaptation to environmental stresses and forms the first defense line against IR. While the repair is a comparatively passive strategy, although it plays a key role in biological evolution.

*D. radiodurans* carotenoids are able to scavenge or quench oxidative and fatal radicals including O_2_^.-^, highly reactive ^.^OH and NO-derived oxidative radicals (Figure 4, 5 and 6). It has been widely accepted that the distinctive behavior of carotenoids is related to their length of the conjugated double bonds and the functional group substitutions of the β-ionone ring [19, 22–24]. As documented in the present study, the two major *D. radiodurans* carotenoid pigments both contained twelve conjugated double bonds with one carbonyl and hydroxyl group (DD-DE) and with two hydroxyl groups (DE) (Figure 1). They might effectively absorb radiation energy or act as direct endogenous radical scavengers, and effectively quench ROS generated during irradiation which can be particularly destructive in biological systems. Apart from its metabolic activities, carotenoid has also been widely noted as a significant structural constituent [26,27]. In the case of *D. radiodurans*, speculation about the possible role of carotenoids in radiation protection has produced similar results [56–58]. Cooperated with a diverse soup of antioxidant metabolites such as pyrroloquinoline quinine (PQQ) [51], natural lipid (vitamin MK8) [52], non-enzymic Mn(II) and Fe(II) ions [3] together make up of the antioxidant/free radical quenching systems in *D. radiodurans*. Therefore, we believe that carotenoids improve the cell’s survival by generically facilitating remove or neutralize ROS generated during irradiation that could cause a series of considerable biological damage to cell structure and functions of all major classes of macromolecules in the cell. Moreover, there are many possible sites where the carotenoids may act.

Intracellular distribution of the components in EDS in *D. radiodurans* has not been fully revealed, although, it’s vital for understanding the function to radioresistance of *D. radiodurans.* Present and forthcoming structural elucidation and cell localization of the unconventional structural components in *D. radiodurans* is not only consequently of significant taxonomic importance, but may also provide new opportunities to evaluate the functions of these components for radioresistance of *D. radiodurans* [41]. The crucial protective evidence of carotenoids in EDS indicates that they make a significant contribution to extraordinary radioresistance of *D*. *radiodurans*. The colorless mutants show substantial sensitivity to γ irradiation, but if these mutations are introduced into the *D. radiodurans* carotenoids, the resulting strains are more resistant to the lethal effects of γ irradiation (Figure 3). Our present study clearly demonstrates potent antioxidant and free radical quenching abilities of two major *D. radiodurans* carotenoids both in vivo and in vitro. Similar observations have published by Tian et al [59] that they evaluated the antioxidant effects of deinoxanthin from *D. radiodurans* R1 using chemiluminescence and DNA damage analyses in vitro, and by Xu et al [60] that they demonstrated the protective effects of carotenoids against radiation and oxidative stress.

Our observations clearly demonstrate that carotenoid is a major natural intracellular component capable of protecting the cells from radiation and oxidative damage by free radical attack during irradiation. The relatively modest effect of carotenoids on cell survival may be due to the fact that carotenoids are not directly involved in latter-stage of DNA repair process when the capacity of EDS can not support the oxidative damage brought by IR to cell structures and macromolecules, but do participate in the earlier-stage of endogenous defense mechanisms of *D. radiodurans* as substantial antioxidants/free radical scavengers at the onset of recovery. Carotenoids, together with other antioxidant enzyme systems and antioxidant metabolites together make up of the EDS in *D. radiodurans*. Since *D*. *radiodurans’*s discovery in 1956, about a half dozen distinct species of the genus has been found out-of-the-way around the world, ranging from animal feces, Mediterranean hot springs, the shielding pond of a radioactive cesium source, the surfaces of Antarctic rocks, in water tanks used as shielding against lethal radiation from pieces of cobalt-60, and even in the feces of some elephants and South American llamas [61–66]. Many studies have shown that many radioresistant bacteria contain red pigments [22,23,62–69]. Because the formation of ROS during irradiation is extremely rapid [7,10], an intracellular protection system which is ubiquitous, but not highly dependent on the induction of enzymes, stage of growth, or temperature over a range at which cells are metabolically active, could provide a selective advantage to the host in oxidative stress environments. Although, there is no direct relationship between bacterial radioresistance and the existence of carotenoid, and currently no available information shown that the carotenoid-pigmented bacteria are more resistant to IR than the other sensitive ones in general, the existence of carotenoids with twelve highly conjugated double bonds and the functional group substitutions of the β-ionone ring in radiation-resistant bacteria but not in sensitive cells support the idea that the presence of carotenoids might be a widespread and active strategy that facilitates survival.

Despite 50 years of investigation, the fundamental questions underlying the extreme resistance phenotype of *D. radiodurans* remain unanswered. Even so, evidence is accumulating that DNA repair strategies could not solely account for its extraordinary radiation resistance of *D. radiodurans*. As discussed in Battista et al [3,5,6], there might exist multiple mechanisms of bacterial radioresistance, including endogenous defense and repair system. Regarding to our investigations reported here, factors may provide a high level of the bacterium radioresistance including the peculiarities of the distinctive cell wall structure, ringlike nucleoids organization, redundancy of genetic information, multiplicity of sites of DNA attachment to the membrane, manganese(II) content, a high level of antioxidant and antiradical systems such as carotenoids. Enhanced radioresistance seems to be the consequence of an evolutionary process that has coordinated various passive and active mechanisms, enabling survival from oxidative stress-inducing conditions such as IR and desiccation [5,6,16]. It is currently difficult to predict which mechanism(s) will be most important for radioresistance, or even whether all of the contributing mechanisms have been discovered. Each of the many enhancements could individually have a modest effect, but collectively they mediate radioresistance. Taken together, our results indicate that carotenoids are crucial to limit the injury produced by an imbalance of deleterious radicals and other ROS, and potential new targets to control recovery from radiation injury. Although we have evaluated antioxidant and radicals quenching effects of carotenoids in *D. radiodurans*, further work is needed to determine the carotenoids biosynthetic genes in *D. radiodurans* and nail down the synergetic effects to understand this bacterium’s unusual talents.

## Materials and methods

### Strains, growth conditions and treatment

Unless otherwise indicated, The wild-type R1 strain of *Deinococcus radiodurans* (ATCC 13939) was used. Cells were incubated in the test medium twice before inoculation. The size of inoculum was 0.1%. All genes were identified as described in the published genome sequence (http://cmr.tigr.org/tigr-scripts/CMR/GenomePage.cgi?org) All strains derived from *D. radiodurans* R1 were grown aerobically at 30 °C in TGY broth (0.5% tryptone, 0.3% yeast extract, and 0.1% glucose, pH 7.0) to an optical density at OD _600_ nm of 0.8-1.0 or on TGY agar (1.5% agar) as described previously [1,4]. *E. coli* strains were grown in Luria-Bertani (LB) broth or on LB plates at 37 °C. At intervals, samples (0.2 ml) were removed from the cultures to determine the cell density. Cell number was determined by direct plate counting. Growth was also monitored photometrically and by dry weight analysis. These methods of biomass measurement resulted in similar growth curves.

*D. radiodurans* cultures in exponential growth were evaluated for their ability to survive DNA-damaging agents. All cultures were treated at 30 °C. The survival was determined by plating serial dilutions of treated cultures in triplicate on TGY plates and incubating at 30°C. The plates were numbered for survivors until 72 h after plating.

Gamma irradiation was conducted using a model 484R ^60^Co irradiator (J. L. Shepherd and Associates, San Fernando, CA, USA) at a rate of 11 Gy/min. Aliquots of the treated cultures in exponential growth were irradiated on ice (0 °C) and removed at one-half-hour intervals over the next 2 h, washed in 10 mM MgSO_4_, and plated on TGY agar to determine cell viability.

### Extraction of *D. radiodurans* carotenoids and chromatographic analysis

The carotenoids in *D. radiodurans* were extracted from 250 mg wet cultures with a 20 ml mixture of cold acetone and methanol (3:2, v/v) supplemented with 0.1% BHT (buthlated hydroxytoluene) as antioxidant, blended in a mechanical blender for 20 minutes, and then centrifuged at 6,000 × *g* at 4 °C for 20 min. The extraction was repeated for three times until all the washing and the residue were devoid of any color The extracts were combined and concentrated under vacuum in a rotary evaporator to dryness in the dark at 30 °C. The extract was transferred quantitatively to a 10 ml volumetric flask using portions of 2 ml of mobile phase for the subsequent LC-MS analysis.

An Agilent polymeric reversed-phase ODS C_18_ column (250 ×4.6 mm i.d.) packed with 5 μm particles (Agilent, CA, USA) and a pump with a diode-array (DAD) detector were used for the HPLC separation and detection. The semi-preparative separation of the carotenoids was applied on a Symmetry Prep™ reverse-phased ODS C_18_ column (7 μm, 7.8 mm × 300 mm I.D.; Waters, Millford, MA, USA), and eluted with a gradient elution consisting of solvent A: 85% acetonitrile, 10% dichloromethane and 5% methanol; solvent B: water with 0-12 min 100% A; and 12-35 min 85% A at a flow rate of 8.0 ml/min using a Waters Delta 600E semi-HPLC equipped with a photodiode array detector from Waters. The purified major product was identified by reverse-phased analytical HPLC with a purity of 97%. LC-MS/MS experiments were performed on a Finnigan LCQ™ Deca XP MAX ion trap mass spectrometer (Thermo Finnigan, CA, USA) by directing the column effluent to an APCI (atmospheric pressure chemical ionization) interface (Finnigan) in the positive mode. The same gradient was used at a flow rate of 0.7 ml/min as mobile phase. Selected [M+H]^+^ was analyzed by collision-induced dissociation (CID) with argon gas, and product ion spectra were recorded.

### Intracellular distribution of *D. radiodurans* carotenoids

The procedure for the preparation of cell walls and membrane was essentially described previously [44]. The cell walls were treated with lytic enzymes and followed by density gradient centrifugation. The individual sediments were extracted and carotenoids were analyzed with the method described above.

### Screening for colorless mutants of *D. radiodurans* by IR

The screening of the mutants with carotenoids synthase genes disruptions in *D. radiodurans* was performed according to the method reported previously [30,60]. For IR, 10^9^ exponential bacterial cells resuspended in 5-ml 10 mM PBS buffer (pH 7.2) were transferred to the glass tube, sealed anaerobically, and irradiated on ice (0 °C) at a rate of 11 Gy/min using a model 484R ^60^Co irradiator (J. L. Shepherd and Associates, San Fernando, CA, USA). The bacterial dilutions after IR were spread on the TGY agar plates. Then all the plates were incubated at 30 °C. The colonies were transferred to nitrocellulose membranes, which provide a white background for visual screening of the clones based on color. Most mutants with different colors were selected until 72 h incubation after plating.

### Assay for antioxidant and free radical scavenging abilities of *D. radiodurans* carotenoids

(1) by UIB system in vivo. The free radical scavenging abilities of antioxidants were determined by measuring the inhibitory rates (%) on UIB (UV-induction of bacteriophage λ) system. The inhibitory rates were calculated according to colony forming unit (CFU) on LB agar plates using the following formula: Inhibitory rate (%) = (CFU_bacterial sample_ - CFU_positive control_) / (CFU_negative control_ - CFU_positive control_) × 100% [36]. (2) by free radical-generation system. The free radical-quenching activities of carotenoids were measured by ultralviolet-visible near-infrared ray (UV/vis/NIR) spectra assay as described previously [70]. Carotenoids were rendered anaerobic by alternating cycles of evacuation and nitrogen purging. The solution was transferred anaerobically to a degassed cuvette sealed with a rubber vaccine cap, such that the cuvette was completely filled leaving no gas phase. Small volume additions of a saturated NO solution in PBS buffer (the NO concentration was calculated to be 12.5 mM using the respective Ostwald coefficient of 0.345) were added to the carotenoid solution to give the molar ratios indicated (carotenoid:NO). The spectra were recorded with a double beam Virian UV-5000 UV-VIS-NIR spectrophotometer (190-1200 nm) (Viran, USA) using 0.1 cm quartz micro-cells. (3) by EPR method. All EPR measurements were performed using a Bruker ESP 300 EPR spectrometer (Bruker Corporation, Germany) operating at X-band with a TM 110 cavity. To measure the generation of free radial, EPR spin-trapping studies were performed using the spin-trapper 5,5-dimethyl-1-pyrroline-N-oxide (DMPO) (Sigma, St. Louis, MO) at 10 mM. EPR spectra were recorded at room temperature in a flat quartz capillary. The instrument setting used in the spin-trapping experiments were as follows: modulation amplitude, 0.95G; time constant 0.2s; scan time 90s; modulation frequency, 25 kHz; microwave power, 10 mW; and microwave frequency, 9.7 GHz. The free radical quenching ability of antioxidant was investigated by comparing EPR spectra in the presence and absence of individual antioxidants.

## Abbreviations

CAT: catalase
CFU: colony forming unit
D_10_: 10% survival value
DD-DE: dehydroxydihydro-deinoxanthin
DE: deinoxanthin
DSB: double-strand breaks
EDS: endogenous defense system
EPR: electron paramagnetic resonance
Gy, grays: H_2_O_2_, hydrogen peroxide
HPI: hexagonal packed intermediate
HPLC: high-pressure liquid chromatography
IR: ionizing radiation
MS: mass spectrum
NO: nitric oxide
O_2_^.−^: superoxide anino/radical
OD_600_: optical density at 600 nm
^.^OH: hydroxyl radical
PQQ: pyrroloquinoline quinine
ROS: reactive oxygen species
SOD: superoxide dismutase
UIB: UV-induction of bacteriophage λ
UV: ultraviolet
UV/vis/NIR: ultraviolet-visible near-infrared ray

## Acknowledgments

We wish to thank Dr. Yiyong Li, Huazhong University of Science and Technology of China, for his excellent assistance with the LC/MS experiment, and Miss Lulu Wang, Wuhan University of China, for the preparation of this manuscript.

## References

1. Anderson AW, Nordon HC, Cain RF, Parrish G, Duggan D (1956) Studies on a radio-resistant micrococcus. I, Isolation, morphology, cultural characteristics, and resistance to gamma radiation. Food Technol 10: 575–578.

2. Brooks BW, Murray RGE (1981) Nomenclature for “*Micrococcus radiodurans*” and other radiation-resistant cocci: *Deinococcaceae* fam. nov. and *Deinococcus* gen. nov., including five species. Int J Syst Bacteriol 31: 353–360.

3. Battista JR (1997) Against all odds: the survival strategies of *Deinococcus radiodurans*. Annu Rev Microbiol 270: 203–220.

4. Daly MJ, Gaidamakova EK, Matrosova VY, Vasilenko A, Zhai M, et al. (2004) Accumulation of Mn(II) in *Deinococcus radiodurans* facilitates gamma-radiation resistance. Science 306: 1025–1028.

5. Battista JR, Earl AM, Park MJ (1999) Why is *Deinococcus radiodurans* so resistant to ionizing radiation? Trends Microbiol 7: 362–365.

6. Cox MM, Battista JR (2005) *Deinococcus radiodurans* - the consummate survivor. Nat Rev Microbiol 3: 882–892.

7. Ghosal D, Omelchenko MV, Gaidamakova EK, Matrosova VY, Vasilenko A, et al. (2005) How radiation kills cells: Survival of *Deinococcus radiodurans* and *Shewanella oneidensis* under oxidative stress. FEMS Microbiol Rev 29: 361–375.

8. Mattimore V, Battista JR (1996) Radioresistance of *Deinococcus radiodurans*: functions necessary to survive ionizing radiation are also necessary to survive prolonged desiccation. J Bacteriol 178: 633–637.

9. White O, Eisen JA, Heidelberg JF, Hickey EK, Peterson JD, et al. (1999) Genome sequence of the radioresistant bacterium *Deinococcus-radiodurans* R1. Science 286: 1571–1517.

10. Daly MJ, Gaidamakova EK, Matrosova VY, Vasilenko A, Zhai M, et al. (2007) Protein oxidation implicated as the primary determinant of bacterial radioresistance. PLoS Biol 5: e92.

11. Levin-Zaidman S, Englander J, Shimoni E, Sharma AK, Minton KW, et al. (2003) Ringlike structure of the *Deinococcus radiodurans* genome: a key to radioresistance? Science 299: 254–256.

12. Zimmerman JM, Battista JR (2005) A ring-like nucleoid is not necessary for radioresistance in the *Deinococcaceae*. BMC Microbiol. 5: 17.

13. Tanaka M, Earl AM, Howell HA, Park MJ, Eisen JA, et al. (2004) Analysis of *Deinococcus radiodurans*’ transcriptional response to ionizing radiation and desiccation reveals novel proteins that contribute to extreme radioresistance. Genetics 168: 21–33.

14. Harris DR, Tanaka M, Saveliev SV, Jolivet E, Earl AM, et al. (2004) Preserving genome integrity: the DdrA protein of *Deinococcus radiodurans* R1. PLoS Biol 2: 1629–1639.

15. Narumi I, Satoh K, Cui S, Funayama T, Kitayama S, et al. (2004) PprA: a novel protein from *Deinococcus radiodurans* that stimulates DNA ligation. Mol Microbiol 54: 278–285.

16. Narumi I (2003) Unlocking radiation resistance mechanisms: Still a long way to go. Trends Microbiol 11: 422–425.

17. Edwards JS, Battista JR (2003) Using DNA microarray data to understand the ionizing radiation resistance of *Deinococcus radiodurans*. Trends Biotechnol 21: 381–382.

18. Liu Y, Zhou J, Omelchenko MV, Beliaev AS, Venkateswaran A, et al. (2003) Transcriptome dynamics of *Deinococcus radiodurans* recovering from ionizing radiation. Proc Natl Acad Sci USA. 100: 4191–4196.

19. Schmidt-Dannert C, Umeno D, Arnold FH (2000) Molecular breeding of carotenoid biosynthetic pathways. Nat Biotechnol 18: 750–753.

20. Albrecht M, Takaichi S, Steiger S, Wang ZY, Sandmann G. (2000) Novel hydroxycarotenoids with improved antioxidative properties produced by gene combination in *Escherichia coli*. Nat Biotechnol 18: 843–846

21. Singh DK, Lippman SM (1998) Cancer chemoprevention—Part 1: Retinoids and carotenoids and other classic antioxidants. Oncology NY 12: 1643–1653.

22. Lemee L, Peuchant E, Clerc M. Brunner M. Pfander H (1997) Deinoxanthin: a new carotenoid isolated from *Deinococcus radiodurans*. Tetrahedron 53: 919–926.

23. Saito T, Ohyama Y, Ide H, Ohta S, Yamamoto O (1998) A carotenoid pigment of the radioresistant bacterium *Deinococcus radiodurans*. Microbios 95: 79–90.

24. Rodriguez-Amaya DB. A guide to carotenoid analysis in foods. International Life Sciences Institute Press, pp 1–60.

25. Britton G (1995) UV/visible spectroscopy. In Britton G, Liaaen-Jensen S, Pfander H (Eds), Carotenoids: spectroscopy, Vol 1B. Birkhäuser Verlag, Basel, pp 13–63.

26. Work E, Griffiths H (1968) Morphology and chemistry of cell walls of *Micrococcus radiodurans*. J Bacteriol 95: 641–657.

27. Hahn M., Saxton WO (1986) Three-dimensional structure of the regular surface layer (HPI layer) of *Micrococcus radiodurans*. J Mol Biol 187: 241–253.

28. Müller DJ, Baumeister W, Engel A (1996) Conformational change of the hexagonally packed intermediate layer of *deinococcus radiodurans* monitored by atomic force microscopy, J Bacteriol 178: 3025–3030.

29. Baumeister W, Karrenberg F, Rachel R, Engel A, Heggeler BT et al. (1982) The major cell envelope protein of *Micrococcus radiodurans* (R1) Eur J Biochem 125: 535–544.

30. Zhang L, Yang Q, Luo XS, Fang CX, Zhang Q, et al (2007) Knockout of crtB or crtI gene blocks the carotenoid biosynthetic pathway in *Deinococcus radiodurans* R1 and influences its resistance to oxidative DNA-damaging agents due to change of free radicals scavenging ability. Arch Microbiol 188: 411–419.

31. Valgimigli L, Pedulli GF, Paolini M (2001) Measurement of oxidative stress by EPR radical-probe technique Free Radic Biol Med 31: 708–716.

32. Ha HC, Sirisoma NS, Kuppusamy P, Zweier JL, Woster PM, et al. (1998) The natural polyamine spermine functions directly as a free radical scavenger. Proc Natl Acad Sci USA 95: 11140–11145.

33. Gao Y, Kispert LD (2003) Reaction of carotenoids and ferric chloride: equilibria, isomerization, and products. J Phys Chem 107: 5333–5338.

34. Furchgott RF, Zawadzki JV (1980) The obligatory role of endothelial cells in the relaxation of arterial smooth muscle by acetylcholine. Nature 288:373–376.

35. Afanas'ev IB. (2005) Superoxide and nitric oxide in pathological conditions associated with iron overload: the effects of antioxidants and chelators. Curr Med Chem 12: 2731–2739.

36. Yang Q, Zhang XL, Zhang JX, Fan CP, Fang CX (2006) A novel method for evaluating free radical scavenging abilities of antioxidants using ultraviolet induction of bacteriophage lambda. J Biochem Biophys Methods 67: 163–171.

37. Fong NJ, Burgess ML, Barrow KD, Glenn DR (2001) Carotenoid accumulation in the psychrotrophic bacterium *Arthrobacter agilis* in response to thermal and salt stress. Appl Microbiol Biotechnol 56: 750–756.

38. Carbonneau MA, Melin AM, Perromat A, Clerc M (1989) The action of free radicals on *Deinococcus radiodurans* carotenoids. Arch Biochem Biophys 15: 244–251.

39. Halliwell B Gutteridge JM (1999) Free Radicals in Biology and Medicine, third ed. Oxford University Press, Oxford.

40. Imlay JA (2003) Pathways of oxidative damage. Annu Rev Microbiol 57: 395–418.

41. Whitehead K, Kish A, Pan M, Kaur A, Reiss DJ, et al. (2006) An integrated systems approach for understanding cellular responses to gamma radiation. Mol Syst Biol 2: 47.

42. Yeaman MR, Yount NY (2003) Mechanisms of antimicrobial peptide action and resistance. Pharmacol Rev 55: 27–55.

43. Kimura M, Anzai H, Yamaguchi I (2001) Microbial toxins in plant-pathogen interactions: Biosynthesis, resistance mechanisms, and significance. J Gen Appl Microbiol 47: 149–160.

44. Adson R and Kansen K (1985) Structure of a novel phosphoglycolipid from *Deinococcus radiodurans.* J Bio Chem 260:12219–12223.

45. Merrick T P, Bruce AK (1965) Radiation Response of Potassium Efflux in *Micrococcus radiodurans* and *Sarcina lutea.* Radiat Res 24: 612–618.

46. Lancy PJr, Murray RGE (1978) The envelope of *Micrococcus radiodurans*: isolation, purification, and preliminary analysis of the wall layers. Can J Microbiol 24: 162–176.

47. Work E, Griffiths H (1968) Morphology and Chemistry of Cell Walls of *Micrococcus radiodurans.* J Bacteriol 95: 641–657.

48. Thompson BG, Murray RGE (1981) Isolation and characterization of the plasma membrane and the outer membrane of *Deinococcus radiodurans* strain Sark. Can J Microbiol 27: 729–734.

49. Meunier-Jamin C, Kapp U, Leonard GA, McSweeney S (2004) The structure of the organic hydroperoxide resistance protein from *Deinococcus radiodurans*. Do conformational changes facilitate recycling of the redox disulfide? J Biol Chem 279: 25830–25837.

50. Wood ZA, Poole LB, Karplus PA (2003) Peroxiredoxin evolution and the regulation of hydrogen peroxide signaling. Science 300: 650–653.

51. Khairnar NP, Misra HS, Apte SK (2003) Pyrroloquinoline–quinone synthesized in *Escherichia coli* by pyrroloquinoline–quinone synthase of *Deinococcus radiodurans* plays a role beyond mineral phosphate solubilization. Biochem Biophys Res Commun 312: 303–308.

52. Reeve J, Kligman LH, Anderson R (1990) Are natural lipids UV-screening agents? Appl Microbiol Biotechnol 33: 161–166.

53. Redpath JL, Patterson LK (1978) The effect of membrane fatty acid composition on the radiosensitivity of *E. coli* k-1060. Radiat Res 75: 443–447.

54. Sistrom WR, Griffiths M, Stannier RY (1956) The biology of a photosynthetie bacterium which lacks colored carotenoids. J Cell Comp Physiol 48: 473–516.

55. Burton GW, Ingold KU (1984) Beta-carotene: an unusual type of lipid antioxidant. Science 224: 569–573.

56. Carbonneau MA, Melin AM, Perromat A, Clerc M (1989) The action of free radicals on Deinococcus radiodurans carotenoids. Arch Biochem Biophys 15: 244–251.

57. Lewis NF, Alur MD, Kumta US (1974) Role of carotenoid pigments in radio-resistant micrococci. Can J Microbiol 20: 455–459.

58. Moseley, B. E. B (1963) The variation in X-ray resistance of *Micrococcus radiodurans* and some of its less-pigmented mutants. Int J Radiat Biol 6: 489.

59. Tian B, Xu Z, Sun Z, Lin J, Hua YJ (2007) Evaluation of the antioxidant effects of carotenoids from *Deinococcus radiodurans* through targeted mutagenesis, chemiluminescence, and DNA damage analyses. Biochim Biophys Acta. 1770: 902–911.

60. Xu ZJ, Tian B. Sun JT, Lin J, Hua YJ (2007) Idenfication and functional analysis of a phytoene desaturase gene from the extremely radioresistant bacterium *D. radiodurans*. Microbiol 153: 1642–1652.

61. Makarova KS, Aravind L, Wolf YI, Tatusov RL, Minton KW, et al. (2001) Genome of the extremely radiation-resistant bacterium deinococcus radiodurans viewed from the perspective of comparative genomics. Microbiol Mol Biol Rev. 65: 44–79.

62. Yoshinaka T, Yano K, Yamaguchi H (1973) Isolation of highly radioresistant bacterium, *Arthrobacter radiotolerans* nov. sp.. Agric Biol Chem 37: 2269–2275.

63. Lewis NF (1971) Studies on radio-resistant coccus isolated from Bombay duck (*Harpodon nehereus*). J Gen Microbiol 66: 29–35.

64. Davis NS, Silverman GJ, Masurovsky EB (1963) Radiation-resistant, pigmented coccus isolated from haddock tissue. J Bacteriol 86: 294–298.

65. Chen MY, Wu SH, Lin GH, Lu CP, Lin YT, et al. (2004) *Rubrobacter taiwanensis* sp. nov., a novel thermophilic, radiation-resistant species isolated from hot springs. Int J Syst Evol Microbiol 54: 1849–1855.

66. Kobatake M, Tanabe S, Hasegawa S (1973) New Micrococcus radioresistant red pigment, isolated from Lama glama feces, and its use as microbiological indicator of radiosterilization. C R Seances Soc Biol Fil 167: 1506–1510.

67. Saito T, Terato H, Yamamoto O (1994) Pigments of *Rubrobacter radiotolerans*. Arch Microbiol 162: 414–421.

68. Galinato MG, Niedzwiedzki D, Deal C, Birge RR, Frank HA (2007) Cation radicals of xanthophylls. Photosynth Res 94: 67–78.

