## Supplemental Figures for "Crucial Roles of Carotenoids as Bacterial Endogenous Defense System for Bacterial Radioresistance of *Deinococcus radiodurans*"

**Supporting information**

**Figure S1.** HPLC analysis showing the intracellular distribution of *D. radiodurans* carotenoids.

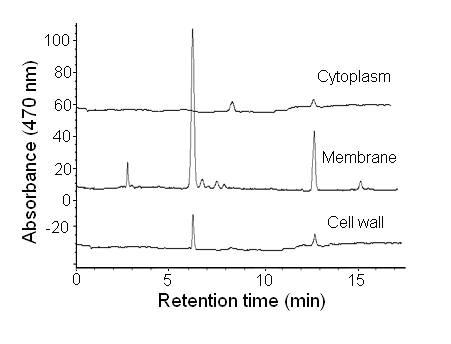

**Figure S2.** HPLC profile of *D. radiodurans* carotenoids isolated from the wild-type and colorless mutants.

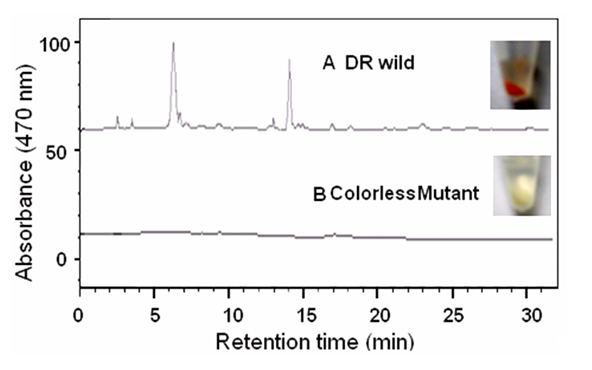
